# The outer-hair-cell *RC* time constant: A feature, not a bug, of the mammalian cochlea

**DOI:** 10.1101/2022.02.02.478769

**Authors:** Alessandro Altoè, Christopher A. Shera

## Abstract

The cochlea of the mammalian inner ear includes an active, hydromechanical amplifier thought to arise via the piezoelectric action of the outer hair cells (OHCs). A classic problem of cochlear biophysics is that the long resistance-capacitance (*RC*) time constant of the hair-cell membrane produces an effective cut-off frequency much lower than that of most audible sounds. The long *RC* time constant implies that the OHC receptor potential—and hence its electromotile response—decreases by several orders of magnitude over the frequency range of hearing. This “*RC* problem” is often invoked to question the role of cycle-by-cycle OHC-based amplification in mammalian hearing. Here, we use published data and simple physical reasoning to show that the *RC* problem is, in practice, a relatively minor physical issue whose importance has been unduly magnified by viewing it through the wrong lens. Indeed, our analysis indicates that the long *RC* time constant is actually beneficial for hearing, reducing noise and distortion while increasing the fidelity of cochlear amplification.

## Introduction

To boost the sensitivity and dynamic range of hearing, the mammalian inner ear embeds a physiologically vulnerable active process that amplifies sound-evoked motions as they propagate as waves along the cochlear spiral (***Shera, 2007***). Although operation of the cellular amplifiers requires prestin-based somatic outer-hair-cell (OHC) electromotility (***Brownell et al., 1985***; ***Zheng et al., 2000***; ***Liberman et al., 2002***; ***Dallos et al., 2008***), serious doubts persist about whether the piezoelectric mechanism can respond with the speed and vigor necessary to boost the power carried by waves of high frequency. This vexing subject has been the focus of much research and debate, starting shortly after the discovery of OHC electromotility (***Santos-Sacchi, 1992***; ***Housley and Ashmore, 1992***) and continuing to the present day (***Vavakou et al., 2019***; ***van der Heijden and Vavakou, 2021***; ***Santos-Sacchi and Tan, 2018***). There are two distinct and largely independent issues. One concerns the kinetics of prestin, which may not be fast enough to support high-frequency electromotility (***Santos-Sacchi and Tan, 2018***).

The second issue, more widely appreciated, arises because OHC electromotility is driven by transmembrane voltage (***Santos-Sacchi and Dilger, 1988***). Electrically, the OHC membrane consists primarily of a resistance (*R*) and a shunt capacitance (*C*) that combine to create a low-pass filter with time constant τ = *RC*. With one exception (***Johnson et al., 2011***), both direct (***Housley and Ashmore, 1992***) and indirect (***Vavakou et al., 2019***) observations indicate that in the basal, high-frequency region of the cochlea, the time constant τ is much larger than the characteristic oscillation period of the basilar-membrane (BM) mechanical response. Equivalently, the cutoff frequency of the OHC low-pass filter is much lower than the maximum frequency of hearing. Consequently, the low-pass filtering is presumed to render the oscillatory component of the OHC receptor potential too small to subserve high-frequency amplification. Dubbed the outer-hair-cell “*RC* time-constant problem” (***Ashmore, 2008***)—or, more compactly, the “*RC* problem” (***Vavakou et al., 2019***)—the dilemma has prompted the search for compensatory mechanisms (***Nobili and Mammano, 1996***; ***Lu et al., 2006***; ***Dallos and Evans, 1995***; ***Ospeck and Iwasa, 2012***) while challenging the biophysical relevance of “cycle-by-cycle” electromotility to cochlear amplification (***Santos-Sacchi, 1992***; ***van der Heijden and Vavakou, 2021***). Ultimately, any definitive evaluation of the significance of these two issues awaits a more complete understanding of OHC dynamics *in situ*, as the mechanical load greatly influences their electromechanical properties (***Ashmore, 1987***; ***Mountain and Hubbard, 1994***; ***Liu and Neely, 2009***; ***Iwasa, 2017***; ***Rabbitt, 2020***).

In this paper we side-step the issue of prestin kinetics and deal solely with the *RC* problem. In effect, we follow classic results about the speed of the OHC “motor” (***Frank et al., 1999***), although these have recently been questioned (***Santos-Sacchi and Tan, 2018***). To bring the RC problem maximally to the fore, we play devil’s advocate and purposely ignore the many proposed avenues for ameliorating the problem. These include the entirely plausible possibilities that (i) dynamic interactions with realistic mechanical loads (***Iwasa, 2017***; ***Rabbitt, 2020***), (ii) the tonotopic variation of OHC electrical properties (***Johnson et al., 2011***), and (iii) the voltage-dependence of OHC baso-lateral currents (***Housley and Ashmore, 1992***; ***Perez-Flores et al., 2020***) may all make beneficial contributions to high-frequency electromotility, both individually and in combination. Thus, we consciously inflate the *RC* problem to its maximal size. Nevertheless, our analysis of this artificially grim, worst-case scenario reveals that, despite its rather long *RC* time constant, the OHC still produces high-frequency (>50 kHz) motile responses comparable to or greater than the motion of the basilar membrane (BM) measured at moderately high sound levels. These results are consistent with recent *in vivo* recordings from the 10 kHz region of the mouse cochlea, which indicate that high-frequency OHC electromotile responses can be large compared to the motions of the surrounding tissue (***Dewey et al., 2021***).

Furthermore, we draw on functional analogies with existing electronic circuits to elucidate multiple ways that the long OHC-membrane time constant appears beneficial for hearing. Among these benefits are significant reductions in the distortion and noise that are the inevitable byproducts of active, nonlinear OHC-based amplification.

### A linearized model of the OHC membrane

To illustrate the ostensible problems that arise from the electrical properties of the OHC, we outline a simple, albeit over-linearized treatment of the generic electrical model of the OHC shown in Fig. 1(a). Our treatment follows that of studies that argue against the significance of cycle-by-cycle OHC amplification. Briefly, mechanically-gated mechanoelectrical transduction (MET) channels drive the electrical response of the OHC basolateral membrane. The electrical impedance of the basolateral membrane is determined primarily by (i) outward K^+^ currents, whose action is collectively represented by a voltage-dependent resistor (***Johnson et al., 2011***); (ii) the membrane capacitance; and (iii) a complex impedance that captures the electrical effects of the mechanical load on the OHC (***Iwasa, 2017***). Motion of the OHC stereocilia modulates the ionic current that flows through the MET channels:

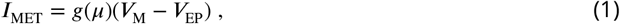

where *μ* is the stereocilia displacement, *g*(*μ*) is the transduction function (units of [Ω]−1), *V*_EP_ the endocochlear potential, and *V*_M_ the OHC membrane potential. For simplicity, we ignore the still unresolved effects of adaptation (***Caprara et al., 2019***) on *μ* and *g*. Accounting for such effects in a linearized model amounts to high-pass filtering the model input and does not modify our conclusions.

**Figure 1.**
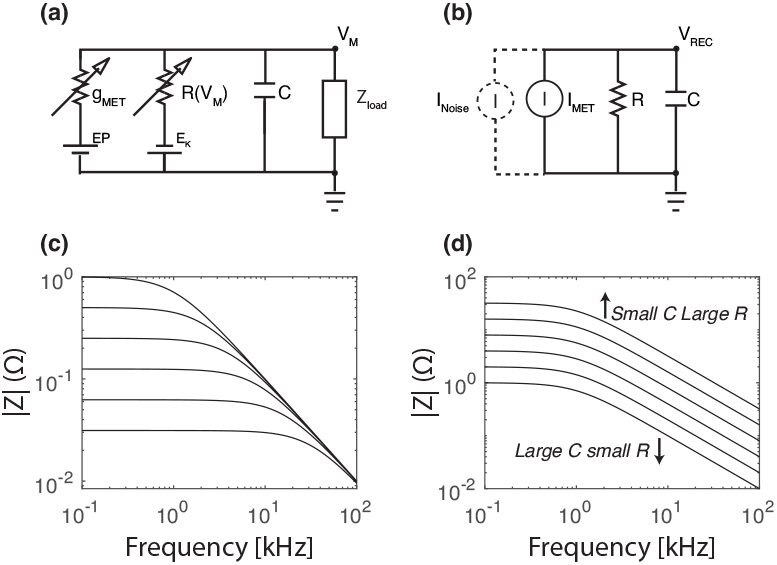
**(a)** Simplified model of the OHC membrane potential. **(b)** Over-linearized model of the OHC receptor potential often used to illustrate the *RC* problem (e.g., ***Vavakou et al., 2019***). **(c,d)** Magnitude of the electrical impedance [*Z*, Eq. (4)] of the *RC* circuit in panel (b). Panel (c) plots *Z* vs frequency for various values of the RC time constant (τ = *RC*) when the capacitance (*C*) is kept fixed; in this case, τ varies linearly with *R*. Conversely, panel (d) plots *Z* when *R* and *C* are covaried with their product, τ, held constant. Together, the two panels demonstrate graphically that at frequencies above the low-pass cut-off frequency, *Z* depends entirely on the capacitance and not on the time constant [Eqs. (4,5)].

Rewriting *V*_M_ as the sum of the resting (*V*_REST_) and receptor (*V*_REC_) potentials yields

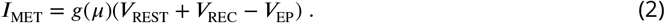

In the small-signal limit (|*V*_REC_| ≪ |*V*_REST_−*V*_EP_|), the dependence of *I*_MET_ on *V*_REC_ can be neglected, and *I*_MET_ becomes proportional to the transduction function, *g*(*μ*). When the stereocilia displacement *μ* is a sinusoid of amplitude *A* and frequency *f*, Taylor expansion of *g*(*μ*) leads to

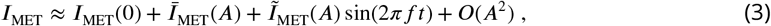

where 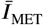 and 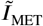 represent, respectively, the DC and AC components of the MET current. The DC component arises from the quadratic term in the series expansion and reflects the possible asymmetry of the transduction function around *μ* = 0.

To highlight the problem of the OHC time constant in its most severe form, we now (over-)linearize the model of the OHC membrane. For example, we ignore the voltage-dependent activation of basolateral currents (***Housley and Ashmore, 1992***; ***Perez-Flores et al., 2020***). When properly linearized, these introduce inductances in parallel with the capacitance, thereby reducing the hair-cell reactance (***Altoè et al., 2018***).^1^ In addition, we ignore the possible salutary effects of the mechanical load [*Z*_load_ in Fig. 1(a)], which may help reduce the effective membrane capacitance *in vivo* (***Iwasa, 2017***). With these simplifications, the OHC membrane appears as the parallel combination of a resistance and capacitance, driven by a (MET) current generator [Fig. 1(b)]. In this worse-case scenario, the electrical impedance of the OHC membrane becomes equivalent to an *RC* (low-pass) filter:

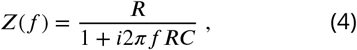

whose associated 3-dB cut-off frequency is *f*_*c*_ = [2*πRC*]^−1^.^2^

For what follows, we find it convenient to introduce a second current generator, in parallel with the MET current, to quantify the effects of various form of noise on the OHC output.

### The time constant *RC* matters less than *R* and *C*

Figure 1(c,d) shows the magnitude of the OHC impedance [Eq. (4)] for various values of *R* and *C*. The resulting curves represent the magnitude of the OHC receptor potential produced by a sinu-soidal MET current of unit amplitude. Figure 1(c) shows the value of the impedance when *C* is fixed and *R* is varied. Although the *RC* time constant decreases (the cut-off frequency increases) with *R*, the magnitude response at high frequencies does not change. Equation (4) implies that decreasing *R* increases the cut-off frequency by reducing the impedance at low frequencies, while leaving it unaltered at high frequencies. (Recall that the cut-off frequency is defined as the frequency where the magnitude falls to 3-dB below its maximum value.) Although one can increase the cut-off frequency arbitrarily by decreasing the response at low frequencies, where |*Z*| is the largest, such manipulations do not improve the high-frequency response—as recently highlighted by ***van der Heijden and Vavakou (2021)***. As a well-known corollary (***van der Heijden and Vavakou, 2021***), “solutions” to the *RC* problem based on mechanisms that effectively lower the value of *R*, do not help with the high-frequency operation of the OHC—they simply disable the OHC at lower frequencies.

Figure 1(d) illustrates the complementary scenario in which the *RC* time constant is held constant while varying the values of *R* and *C*. Although the cut-off frequency remains fixed, the response amplitude changes dramatically at all frequencies, depending on the relative value of *R* and *C*. In this scenario, |*Z*| increases in proportion to *R* or, equivalently, inversely with *C*. These examples make clear that simply knowing the value of the *RC* time constant reveals little about the magnitude of the OHC response at high frequencies. This observation has important consequences for OHC biophysics: In OHCs, the membrane capacitance *C* is small (a few pF) and the resistance *R* is large (dozens of MΩ).

OHCs in the high-frequency regions of the cochlea operate well above the cut-off imposed by their *RC* time constant. In this case, the current flowing through the resistance *R* becomes small compared to that flowing through the capacitor *C*. Therefore, the AC receptor potential becomes

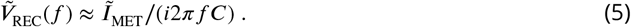

Equivalently, the MET current required to achieve a given “target” value 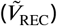 of the OHC receptor is

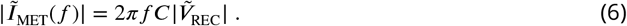

In other words, the so-called “*RC* problem” that supposedly limits the high-frequency operation of the OHC reduces to the problem of charging a capacitor to a certain potential: the *C* problem (© ***van der Heijden and Vavakou, 2021***).

### Little seems enough: Worst-case parameters yield realistic displacements

To explore whether the membrane capacitance limits the high-frequency operation of the OHC, we apply Eq. (5) to estimate the magnitude of OHC electromotility at the highest frequencies for which systematic mechanical recordings of BM and organ-of-Corti motions are available [namely, in the ∼50 kHz region of the mouse cochlea (***Ren et al., 2016b***)]. Again, our focus on the *RC* problem allows us to ignore possible limits imposed by prestin dynamics. Additionally, we side-step the many unknowns concerning OHC force production by comparing estimated limits on OHC length-changes with BM displacements recorded *in vivo*. The argument goes as follows: if the electrical drive to the OHC can produce motile responses comparable to or larger than the measured motion of the BM (and surrounding structures), then the AC response of the OHC is evidently “large enough,” and membrane filtering does not preclude functionally relevant high-frequency operation.

Again, we suppress any bias towards wishful thinking by perversely imagining a worst-case scenario. Consider, for example, the well-known tonotopic variation in OHC properties. Whereas OHC length and capacitance increase systematically along the cochlear spiral, the MET-channel conductance decreases with position (***Beurg et al., 2006***; ***Johnson et al., 2011***; ***Soons et al., 2015***). For a given AC receptor potential, long OHCs elongate more than short ones (***Frank et al., 1999***). On the other hand, short cells have smaller capacitance, and are therefore more easily charged. The MET conductance determines the amplitude of the MET current, and hence the magnitude of the OHC electromotile response. Since our goal is to make unfavorable parameter choices, we take our hypothetical murine OHCs at the 50-kHz place to be simultaneously (i) short and therefore limited in their ability to elongate and (ii) endowed with membrane capacitance and MET currents representative of OHCs from more apical locations with a much lower CF (***Johnson et al., 2011***). In a nutshell, we not only neglect the known tonotopic variations, we purposely counteract them to inflate the apparent severity of the *RC* problem.

Based on the micro-chamber experiments of ***Frank et al. (1999)***, we assume that short OHCs expand and contract by ∼1 nm/mV at frequencies close to 48 kHz. Although the voltage-displacement relation depends on the OHC’s mechanical load—which may not be the same in the micro-chamber as it is *in situ*—this estimate is in good agreement with *in situ* values obtained using electrically evoked responses in the excised temporal bone of the guinea pig (see Fig. 2 of ***Nowotny and Gummer, 2006***). We also assume that the available AC MET current is on the order of 1.5 nA—half of the available MET current measured in gerbil OHCs at the 10-kHz place (***Johnson et al., 2011***)—and assume that the OHC capacitance is ∼4 pF, also based on data from the 10-kHz place in gerbil (***Johnson et al., 2011***).

**Figure 2.**
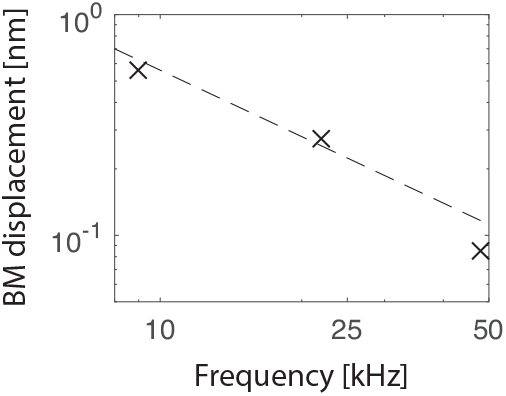
Scaling of BM displacement in the mouse cochlea. The symbols give BM displacement measured using CF tones of 20 dB SPL (where BM responses are approximately linear) at CFs of 9 kHz (***Dewey et al., 2019***), 22 kHz (unpublished data courtesy of J. B. Dewey), and 48 kHz (***Ren et al., 2016b***). The trend is well captured by a function inversely proportional to frequency (dashed line).

Using these values, we find [Eq. (5)] that our worst-case OHC is capable of elongating on the order of 1.25 nm at 48 kHz. For comparison, vibrational data from the 48-kHz region of the mouse cochlea (***Ren et al., 2016b***) indicate that *in vivo* mechanical movements are close to this value at sound levels of 60–70 dB dB SPL (at 48 kHz, BM and RL responses at 70 dB SPL are less than 3 nm). In these experiments, both BM and RL gain at CF decrease with level above 30 dB SPL (where BM motion is ∼0.2 nm) and are dramatically reduced above about 70 dB. That is, our approximate numerical bound on the magnitude of the OHC electromotile response (1.25 nm) is much larger than the BM displacements measured at low sound levels, where amplification is strongest, and is comparable to BM displacement at moderately high levels, where amplification is greatly reduced. Note that if this comparison had come out differently—if our approximate upper bound on OHC electromotility had been much less than the displacement of the BM and surrounding structures—then assertions that cycle-by-cycle somatic motility appears too weak to influence intracochlear motions at high frequencies would be intuitively compelling. As it turns out, however— and notwithstanding a number of unfavorable assumptions—the comparison demonstrates that the low-pass filtering of the OHC membrane does not, in practice, impose intrinsic limitations on the cell’s high-frequency response that compromise its ability to make significant contributions to measured cochlear amplification.

For a complementary perspective, we can run the analysis “in reverse” using Eq. (6). Although it is not easy to determine, on theoretical grounds, how much the OHCs need to elongate and contract in order to provide the required wave amplification, we can infer the voltage and current required to drive OHC motile responses comparable in magnitude to the BM vibrations observed at sound levels near the threshold of hearing. Linear extrapolation of the reference data (***Ren et al., 2016b***) indicates that a 48-kHz tone at 10-dB SPL—a sound level somewhat smaller than the threshold of the most sensitive auditory-nerve fibers (***Taberner and Liberman, 2005***)—evokes BM responses of ∼20 pm, requiring voltage swings on the order of 20 µV, and corresponding MET currents on the order of 25 pA. This current is on the order of the AC current mediated by one or two MET channels.^3^ Recent *in vivo* recordings from the 10-kHz region of the mouse (***Dewey et al., 2021***) indicate that OHC electromotile responses are approximately 4 times larger than BM vibrations near CF. Assuming that this relationship also holds at higher CFs, the simple model implies that OHC near-CF AC potentials in the 50 kHz region must be ∼80 µV near threshold, requiring a driving current equivalent to that mediated by roughly 4–8 MET channels.

Following ***van der Heijden and Vavakou (2021)***, we note that BM nonlinearities and amplification are observed at frequencies where BM (and RL) responses are ∼20 dB smaller than at CF (***Ren et al., 2016b***). At these frequencies, the simple OHC model implies, conservatively, that AC receptor potentials on the order of few (2–8) µV are sufficient to elicit OHC displacements comparable to or greater than the near-threshold motion of the BM.

These values appear small, although they are somewhat larger than the receptor potentials (<1μV) extrapolated by ***van der Heijden and Vavakou (2021)*** from *in vivo* intracellular recordings at the 16 kHz region of the guinea pig (***Cody and Russell, 1987***). Note, however, that the validity of this extrapolation is unclear. It has long been known that the perforation produced by the microelectrode *in vivo* has a major impact on hair-cell electrodynamics (***Kros and Crawford, 1990***), with the likely result being an underestimate of the true physiological range of AC receptor potentials (see ***He et al., 2004***). Furthermore, the invasive approach adopted in such experiments likely results in reduced cochlear sensitivity. Indeed, ***Cody and Russell (1987)*** report that in their experiments, OHC responses (AC, near-CF) at sound levels near neural threshold are roughly 10–30 μV, a range of values that compares favorably to the results presented above.

### The long *RC* time constant reduces noise where it matters most

Do such small voltage excursions and the resulting motile responses have any functional relevance in the presence of noise? Multiple noise sources contribute to the OHC transmembrane voltage, including various forms of hydro-mechanical noise that influence the motion of the stereocilia (***Sasmal and Grosh, 2018***). Although the magnitudes of these noise sources are not yet fully characterized, their effects can be explored in the model by introducing a current source whose output represents the sum of the various forms of white (flat power spectral density), pink (1∕*f*), and brown (1∕*f*^2^) noise expected to be present in the cochlea. These noise sources include, but are not limited to, the thermal noise associated with the OHC membrane resistance, pink noise associated with the kinetics of K+ transport, stochastic gating of the MET channels, and mechanical and background noise contributing to stereocilia displacement.

Note that the OHC membrane filters these noise sources in the same way that it filters the driving MET current, a quantity otherwise known as the “signal” (Fig. 1c). Consequently, the signal-to-noise ratio (SNR) measured in small frequency bands around the stimulus frequency mirrors the ratio of the stimulus-driven MET current to the noise current. Simply put, at any given frequency, the signal-to-noise ratio at the OHC output *is independent of the membrane filtering*.

Details of the power spectral density of the inherent electrical noise (i.e., thermal noise and the flickering of MET channels) depend on OHC parameters (e.g., the fraction of MET channels open at rest) for which there is as yet little agreement in the literature. In Appendix 1, we estimate that the inherent electrical noise corresponds to a broadband current source whose root-mean-square (RMS) amplitude is on the order of that mediated by 1–5 MET channels. Interestingly, this range of values closely matches that required to elicit OHC motions near the threshold of hearing (see above). But note that there are generally three OHCs at each CF location. Simple mathematical reasoning (coherent signal, incoherent noise) implies that if each of the three contributes equally, this architecture could boost the signal-to-noise ratio of the cochlear amplifier by roughly 5 dB. Additionally, whereas tonal stimuli have narrow bandwidths, the noise power of the current source is uniformly distributed over a broad frequency range. Thus, the inherent electrical noise at the hair-cell input—and, consequently, the output—is therefore relatively small.

Importantly, the long *RC* time constant means that noise at the OHC output is significantly attenuated at high frequencies. Indeed, low-pass filtering by the membrane not only reduces the overall noise power at the OHC output but also confines it to a region where it cannot do much harm. According to prevailing theory (e.g., ***Shera, 2007***), the OHCs amplify traveling waves by locally pumping small amounts of power into the adjacent fluids. The coherently emitted^4^ power accumulates spatially and boosts the amplitude of the traveling pressure wave and thereby the motion of the BM. Coherent wave amplification is most effective in the so-called “short-wave region” that extends roughly 0.5–1 octave basal to CF, depending on species and tonotopic location (see ***Altoè and Shera, 2020b***, and Fig. 3d).^5^ Indeed, *in vivo* measurements demonstrate that OHC electrical activity in the gerbil (***Dong and Olson, 2013***) and OHC mechanical activity in the mouse (***Dewey et al., 2019***) are effectively coupled to the traveling wave only in frequency ranges spanning about 0.5 and 1 octave below CF, in excellent agreement with the theory. Furthermore, electrically-evoked mechanical responses *in situ* appear broadband near the apical surface of OHCs but narrowband on the BM (***Ren et al., 2016a***).

**Figure 3.**
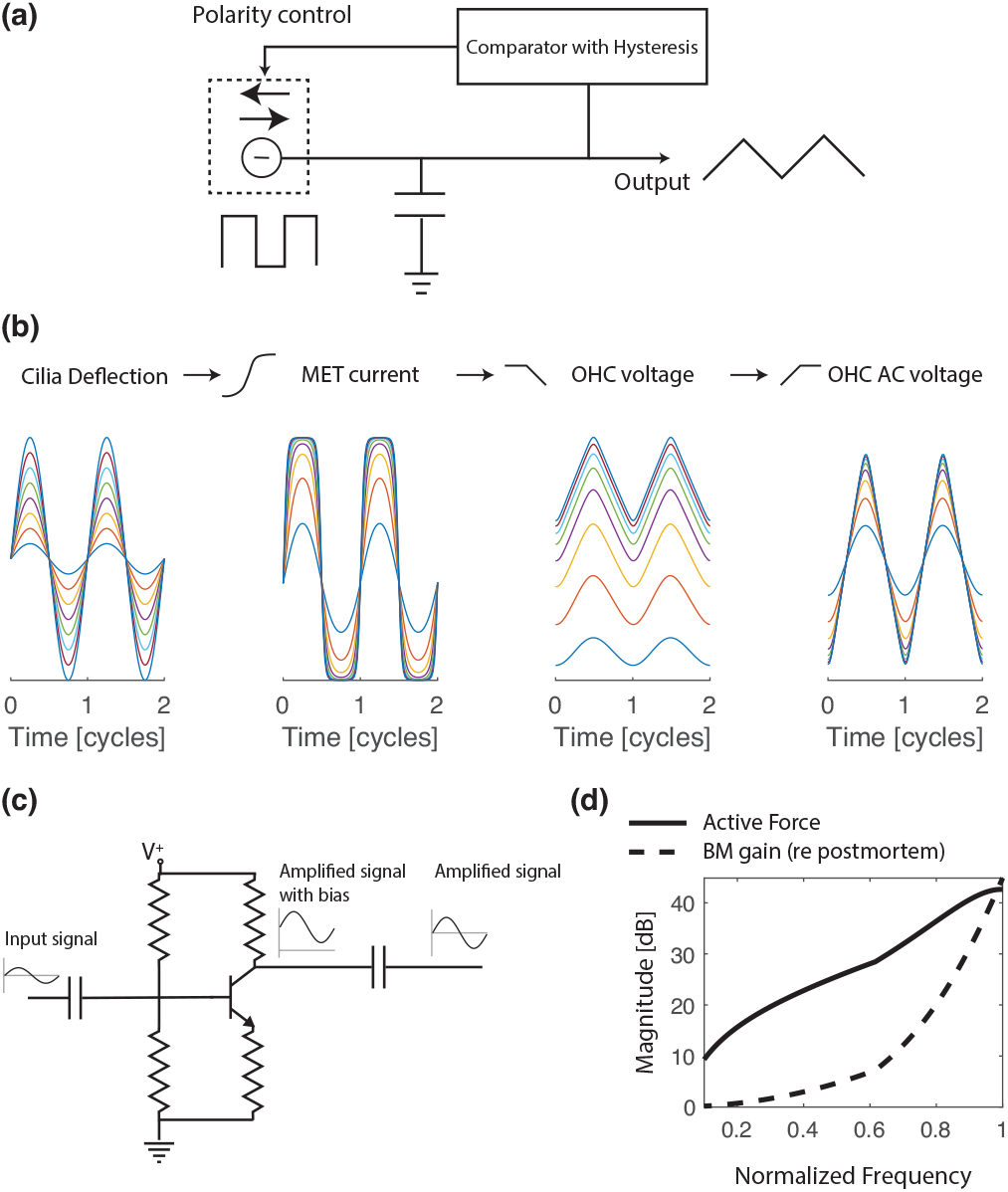
**(a)** Schematic illustration of a current-controlled oscillator, whose operation has important analogies with the OHC [see text and panel (b)]. **(b)** Simple model elucidating the effect of OHC low-pass filtering for improving the fidelity of the hearing organ. From left to right: The input is assumed to be a sinusoidal deflection of the OHC stereocilia. This vibration is converted into MET current via an asymmetric, saturating sigmoidal nonlinearity. The OHC voltage is derived from the MET current by low-pass filtering, in accordance with the model adopted for our analysis of the *RC* problem. The oscillating (AC) component of the OHC voltage is obtained by high-pass filtering the OHC voltage. Whereas the MET current becomes highly saturated and distorted as the stereociliary input increases, the OHC voltage remains much less distorted, and the AC response is nearly sinusoidal. This same model has recently been shown to capture the mechanical distortions measured near the OHCs in the organ of Corti (***Dewey et al., 2021***). **(c)** Example of AC coupling in a textbook signal amplifier. The signal at the output terminal (collector) of the transistor is the sum of the amplified version of the input signal and a DC bias. The series capacitor at the output terminal acts as a high-pass filter, removing the DC bias from the amplified signal. The capacitance value is selected to couple the amplified signal in a relevant range of frequencies. **(d)** In analogy with the signal amplifier of panel (b), OHC forces are AC coupled to the traveling wave on the BM. This panel compares the magnitude of OHC forces and amplification of BM response, quantified as the ratio between BM displacement in both the presence and absence of active forces (i.e., postmortem) in a 3-D model of the mouse cochlea (***Altoè and Shera, 2020b***). Whereas OHC forces are broadband, BM amplification retains a prominent high-pass characteristic, so that low-frequency OHC activity has a negligible effect on the BM response. In this simple model, OHC forces are active (i.e., amplifying and in phase with BM velocity) at all frequencies; the prominent frequency dependence of BM amplification is brought about by the short-wave hydrodynamics.

In a nutshell, the OHC low-pass filter attenuates the OHC response to noise at frequencies where it would otherwise couple strongly to the traveling wave. By decoupling OHC noise from the hydromechanical action of the cochlear amplifier, the long *RC* time constant renders the noise band different from the amplification band. Additionally, present understanding suggets that the inner-hair-cell (IHC) stereocilia are driven primarily by fluid velocity, which is inevitably high-pass filtered relative to the displacement of the surrounding tissue (for a review, see ***Guinan, 2012***). Thus, the long *RC* time constant reduces noise at frequencies where it would interfere most with the operation of the primary sensory cells, at least in the base of the cochlea.

### Tonotopic variations, scaling, and operation at higher frequencies

What about operation at even higher frequencies? Simple arguments suggest that our results, derived from data at ∼50 kHz, extend further. With fixed parameters, the model predicts that the OHC motile response decreases in a manner inversely proportional to frequency. But the same is true both for the expected displacement of the sensory tissue and for the noise at the OHC output (see above). Figure 2 demonstrates that the logarithm of near-CF BM displacement in the mouse decreases almost linearly with frequency.^6^ That is, the estimated frequency-dependence of the OHC response-limit mirrors that of BM displacement.

In the real cochlea, OHC properties vary systematically along the tonotopic axis. Nevertheless, many prominent variations in cochlear responses (such as the change in tuning with CF) can be reasonably accounted for by models where the action of the OHCs is assumed scaling symmetric and thus essentially identical at all locations (***Altoè and Shera, 2020a***), with possible deviations at the extreme apex near the helicotrema (***Sasmal and Grosh, 2019***). We note that a principal functional consequence of the tonotopic gradient of OHC length (***Soons et al., 2015***)—reduced OHC motion at higher CFs—is effectively counteracted by the gradients of OHC capacitance and MET current (***He et al., 2004***; ***Johnson et al., 2011***), which act to increase OHC motion at higher CFs. Thus, an assumed scaling symmetric (or “one-size-fits-all”) phenomenology of OHC action is perhaps not unreasonable. Note also that systematic variations of MET-channel sensitivity (***He et al., 2004***) and cytoarchitectural geometry (***Soons et al., 2015***) may serve to help equalize OHC functionality along the tonotopic axis, and not necessarily differentiate it, as often assumed.

As a corollary, note that the assumption of approximate functional invariance of the OHCs along the cochlea implies that near-CF OHC velocity, like the peak velocity of the BM (see Fig. 2), is independent of location. Because the mechanics of the tissue surrounding the OHC are likely to be viscous-dominated [rather than stiffness-dominated, as recently assumed by ***Vavakou et al., 2019***] at frequencies extending several octaves below CF (***Scherer and Gummer, 2004***), the force that OHCs deliver to the sensory epithelium (and consequently the power injected to the traveling wave) is also approximately independent of CF. This is precisely what active, scaling-symmetric models (e.g., ***Zweig, 1991***) have implemented since the dawn of cochlear amplification (***Kemp, 1978***).

### The long *RC* time constant reduces distortion

We now employ analogies with existing electronic circuits to illustrate how much the *RC* problem has been magnified over the years by an excessive focus on the mental construct of the low-pass filter. We first turn attention to a basic building block of analog electronics: the triangle-core oscillator, a simple circuit whose somewhat abstracted representation is shown in Fig. 3a. The operation of this oscillator is straightforward. A constant positive current charges a capacitor. When the capacitor voltage reaches a desired maximum value, an auxiliary circuit inverts the current polarity, discharging the capacitor over time. When the capacitor voltage reaches a desired minimum value, the current polarity is again inverted, back to positive, and the capacitor is recharged. And so on. The voltage output of the circuit is a triangular waveform whose fundamental frequency depends on the strength of the current source, the capacitance to be charged, and the desired voltage swing. Although theoretically infinite, the time constant of the charging element (the capacitor) is limited in practice by parasitic conductances in the capacitor and by other non-ideal circuit components. This simple circuit provides a concrete, real-life example of a system whose high-frequency operation, like that of the more exotic OHCs, is not limited by the *RC* time constant of its charging element. Indeed, like the OHCs, this circuit operates above the cut-off frequency (theoretically zero) imposed by its *RC* time constant. But perhaps the analogies do not end here.

For example, regarding the OHC as a filter driven by a sinusoidal input may overlook important aspects of OHC function. It is now a classic result that the MET transduction function is a highly nonlinear and saturating function (e.g., ***Russell et al., 1986***; ***He et al., 2004***), implying that the MET current approaches a square wave for sufficiently large excursions of the stereocilia. When the *RC* time constant is much larger than the period of the local CF, the response remains “clean” despite the highly distorted input (Fig. 3b). Thus, in the same way that the capacitor in the electronic oscillator transforms a square current into a triangular voltage, the OHC low-pass filter increases the fidelity of the hearing organ by reducing harmonic distortion. Indeed, a recent study (***Peterson and Heil, 2020***) proposed an analogous role for the low-pass filtering present in the IHC mechano-to-neural transduction machinery. Just as in our simple OHC model [Fig. 3b], IHC low-pass filtering reduces the distortion introduced by the saturating MET nonlinearity, thereby increasing the fidelity and temporal precision of auditory-nerve responses (***Peterson and Heil, 2020***).

### OHCs as regulators?

The analogy with electronic circuits allows one to analyze, with minimal complexity, the recent proposal that the OHCs serve primarily as sluggish regulators rather than as the motors for cycle-by-cycle power amplification (***Vavakou et al., 2019***). The hypothesis is that because their response near DC is large and nonlinear, OHCs act as envelope detectors whose role in cochlear mechanics must be that of somehow modulating the traveling wave by serving as automatic gain controllers (AGCs). Although AGC theories of cochlear mechanics have been criticized and found wanting on other grounds (***Altoè et al., 2017***), a simple analogy with AC- and DC-coupled electronic circuits raises additional theoretical and empirical concerns.

Very low frequency (near-DC) biases introduced by electronic circuits in the signal path are often undesirable (e.g., in radio and audio applications). The problem is commonly solved by connecting a capacitor (or a transformer) in series with the output (or input). The capacitor acts as a high-pass filter, removing the DC component of the signal (Fig. 3c). Using this simple but effective strategy, multiple circuits can be chained together without worrying about the potentially deleterious side-effects of DC biases (or low-frequency noise) in the signal path. Circuits that include these coupling capacitors (e.g., radio receivers and the inputs and outputs of most audio signal processors) are said to be AC-coupled. Conversely, circuits that are designed to generate or process DC or very low-frequency signals (e.g., on/off thermostats) exclude coupling capacitors and are said to be DC-coupled. Basic physics implies that in the mammalian cochlea both slow and compressional pressure waves are AC coupled to the motions of the sensory epithelium (***Peterson and Bogert, 1950***; ***Lighthill, 1981***; ***Shera and Zweig, 1992***). Additionally, OHC activity is theoretically (Fig. 3d) and empirically found to couple to the traveling wave predominantly in a narrowband region below CF (see ***Dong and Olson, 2013***; ***Dewey et al., 2019***, and “*The long RC time constant reduces noise where it matters most*”).

Thus, OHC motions appear strongly AC coupled to the traveling wave, the primary means of energy transport in the cochlea. Forms of DC coupling between OHC motions and the traveling wave—necessarily more indirect than the known AC coupling—have not been identified and proposed models are at odds with the experimental data (***Altoè et al., 2017***). Indeed, theories that propose a central, regulating role for the DC response of the OHC are fundamentally similar to the “impedance-reduction” hypothesis of ***Kolston et al. (1990)***. This hypothesis posits that the OHCs act to regulate the reactive and/or resistive impedance components of the cochlear partition (i.e., its stiffness and/or damping), without boosting the energy carried by the traveling wave (see ***Allen, 2001***; ***van der Heijden and Vavakou, 2021***). Kolston himself ultimately determined that impedance-reduction models do not fit the data (***Kolston, 2000***). In particular, Kolston demonstrated that impedance-reduction models tailored to replicate the BM-magnitude response predict excessive phase accumulation, incompatible with the data. He concluded that the measured BM phase response implies some form of energetic boost to the BM wave. As it turns out, this result is no mere model-dependent speculation: the real and imaginary parts of the wavenumber—which determine the phase and energy accumulation, respectively—are not independent of one another, but obey dispersions relations imposed by causality (see ***Shera, 2007***). Furthermore, we have recently demonstrated (***Altoè and Shera, 2020b***) that significant level-dependencies of the partition damping and stiffness cannot easily be reconciled with the approximate level-invariance of the fine-time structure of the BM click-response and the biological robustness of this phenomenon (for a review, see ***Shera, 2001***; ***Zweig, 2016***). Indeed, the problem would be particularly acute were the level-dependence to mirror OHC activity and occur in extended regions basal to the peak of the traveling wave (see Appendix A of ***Altoè and Shera, 2020b***).

The results of our simple model (see “*The long RC time constant reduces noise where it matters most*”), taken together with the inevitability of noise sources having 1∕*f* (or 1∕*f*^2^) spectral densities, leave little doubt that the noise at the OHC output must be substantially larger at low (near-DC) frequencies than at high. When the noise sources are white, the SNR at the OHC output measured in narrow frequency intervals for a given oscillatory MET current is fairly independent of frequency. But in the real world, where many noise sources are pink and brown, the SNR is expected to decrease significantly at low frequencies. Unfortunately, providing more quantitative statements about the effects of noise in alternative “gain-control” scenarios remains difficult in the absence of concrete, mechanistic proposals. However, physical intuition suggests that the low-pass nature of physical and biological noise can only render noise more problematic for mechanisms based on the DC response of the OHC (***Vavakou et al., 2019***; ***van der Heijden and Vavakou, 2021***) than for existing theories where OHCs are AC coupled to the traveling wave.

### The *RC* problem and OHC tuning

A classic problem associated with the OHC membrane circuitry is that low-pass filtering is a nuisance when attempting to produce sharply tuned OHC forces. Although solutions to this problem have been proposed (e.g., ***Lu et al., 2006***), we argue that the 3-D nature of cochlear hydrodynamics renders the problem moot. To compensate for the oversimplified hydrodynamics (***Shera et al., 2005***), sharply tuned OHC forces acting on the BM are indeed necessary in cochlear models with zero-or one-dimensional fluid coupling. (Examples of zero- and one-dimensional models include filter-bank/critical-oscillator and transmission-line models, respectively.) However, models that incorporate two- or three-dimensional fluid coupling can produce realistic narrowband BM amplification without resorting to sharply tuned OHC forces. This is nicely illustrated by a recent 3-D model that replicates BM data well—and provides intuitive explanations for a number of other observations—while employing phenomenological OHC forces that are low-pass filtered relative to the BM (see ***Altoè and Shera, 2020b***, and Fig. 3d). Similar results are found in the data. Indeed, a recent study in the ∼9 kHz region of the mouse (***Dewey et al., 2021***) demonstrates that the OHC motile response *in vivo* appears low-pass filtered relative to that of the BM; the cut-off frequency associated with such low-pass filtering is suspiciously similar to OHC corner frequencies reported in the literature. Despite the prominent low-pass filtering, the cycle-by-cycle OHC response is larger than that of the BM, even at CF and at stimulus levels of 70 dB SPL, corroborating the analysis presented above, as well as its conservative character.

We conclude by noting what is now obvious from the literature: detailed, physics-based cochlear models that include OHCs with membrane time constants similar to those reported in the literature have no problem producing realistic wave amplification (e.g., ***Sasmal and Grosh, 2019***). Arguments suggesting that the low-pass filtering action of the OHC membrane precludes effective cycle-by-cycle wave amplification in the cochlea evidently fall prey to the fallacy of composition— mistaking the parts (an isolated OHC) for the whole (the cochlear amplifier). For example, one can easily obtain high-, band-, and low-pass transfer functions by cascading multiple low-pass filters and mixing their responses, with or without the introduction of feedback loops.^7^ In a similar way, the cochlear amplifier can be thought of as emerging from the “mixed” responses of multiple OHCs. Mixing occurs hydrodynamically, via the fluids and other coupling mechanisms, and each coupling mechanism has its own frequency (or wavelength) dependencies. Thus, it should not be surprising that the frequency-dependence of cochlear amplification differs from that of the isolated OHC. Indeed, while OHC responses are low-pass filtered relative to that of the BM, they are band-pass relative to the cochlear input [whether that input is considered to be the pressure near the tympanic membrane or the vibration of the stapes (***Lee et al., 2016***; ***Ren et al., 2016b***; ***Dewey et al., 2021***)], a direct consequence of the tuned mechanical drive to the OHC stereocilia. Despite a mischievious grin that has taunted the field for decades, the *RC* problem, like the Cheshire cat, appears effectively to have vanished from view in both modern theories and experiments.

## Acknowledgments

This work was supported by NIH Grant R01 DC003687 (CAS).

## Appendix

Appendix 1

Estimate of electrical noise

Electrical noise is conveniently modeled as a current generator driving the OHC membrane (Fig. 1b). The equivalent thermal noise current source has power spectral density

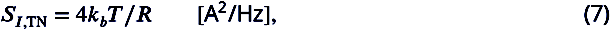

where *k*_*b*_ is the Boltzmann constant, *T* the absolute temperature and *R* the OHC resistance. Note that this current generates a voltage at the OHC output with RMS amplitude 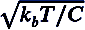 and bandwidth 1∕4*RC*. At body temperature, and assuming a resistance *R* of approximately 10–20 MΩ, Eq. (7) indicates the noise current power in the frequency band of 0–50 kHz is the same as that produced by a sinusoidal current of ∼10 pA, which is on the order of that mediated by a single MET channel (***Beurg et al., 2006***).

The “shot” noise (***van der Heijden and Vavakou, 2021***) associated with the random opening and closing of MET channels is not so easy to estimate. Regardless of the assumed stochastic behavior of MET-channel gating, the shot-noise spectral density—with units of A^2^/Hz—is proportional to *qI*_MET_, where *q* is the charge mediated by a single opening event. Unfortunately, *q* is hard to estimate from published experiments without making a number of assumptions. If the MET channels open and close independently of one another, the root-mean-square (RMS) noise current is on the order of

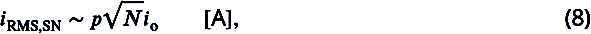

where *p* is the channel open-probability, *N* the number of MET channels, and *i*_0_ the current mediated by a single channel. Although the value *p* “at rest” remains controversial— *in vitro* estimates range from roughly from 10–50% (***Russell et al., 1986***; ***Johnson et al., 2011***)—relatively strong rectification and even-order harmonics are observed in mechanical responses from the OHC region in both the base and apex of the mouse (***Dewey et al., 2021***; ***He and Ren, 2021***), and in the base of the gerbil as well (***Vavakou et al., 2019***), suggesting that *p* is relatively small (i.e., ≤ 20%). The number of MET channels is on the order of 100 [∼60 tip-links, with on average of 1.65 channels per tip-link (***Beurg et al., 2006***)]. With these values, Eq. (8) implies an RMS shot-noise similar to the current mediated by a small number of channels [0–5 for *p* ∈ (0, 0.5)].

As an interesting aside, note that this simple analysis points out that the degree of rectification (i.e., the value of *p* at rest, or equivalently the asymmetry in the MET transduction function) controls the OHC susceptibility to shot noise, with smaller values of *p* reducing the noise at the OHC input and, consequently, at the OHC output. On the other hand, in the absence of noise, the OHC is maximally sensitive to small stereocilia vibrations when *p* = 0.5 and the transduction nonlinearity is perfectly symmetric (***Johnson et al., 2011***). Therefore, the asymmetry of the MET transduction function (i.e., the value of *p*)—and its possible systematic tonotopic variation (***Nam and Guinan, 2016***)—may play an important role in controlling the effective OHC sensitivity in the presence of noise. Although *p* = 0.5 optimizes signal detection in idealized, noiseless models, the same is not true in the presence of shot noise.

## Footnotes

1 Because it ignores leading-order linear terms that emerge when properly linearizing the basolateral K^+^ currents (see ***Koch, 2004***), this “straightforward linearization” (***Vavakou et al., 2019***) is not a true linearization of the OHC membrane model. Nonetheless, the simplification appears to capture the OHC motile response *in vivo* reasonably well (***Vavakou et al., 2019***; ***Dewey et al., 2021***).

2 In the literature the resting MET conductance is sometimes included in the value of *R* (e.g., ***Johnson et al., 2011***). The definition of the *RC* time constant requires a linearized model, and linearization when the input of the model is the vibration of the sterocilia leads to representing the MET channels as a current generator [Fig. 1(b)]. We therefore find it more informative to exclude the MET resting conductance from the calculation of *R*. Indeed, while the MET resting current indirectly contributes to the filtering property of the OHC (by determining resting potential and hence the resting basolateral conductance), once this conductance is determined and “fixed” in the model, the oscillating (AC) component of the OHC potential is independent of the MET resting current.

3 ***Beurg et al. (2006)*** report that a single MET channel in the mid-turn of the rat cochlea mediates a current of ∼12 pA and that the channel conductance increases with CF. If there are approximately 100 MET channels per OHC (***Beurg et al., 2006***), then a single channel in a 10-kHZ gerbil cell mediates a current of ∼3 nA∕100 = 30 pA (***Johnson et al., 2011***). Not knowing the single-channel current in the 50-kHz region of the mouse cochlea, we use these two values as proxies.

4 A prominent feature of theories of cochlear wave amplification is that the amplification of spatially distributed intracochlear sources, such as the motions produced by OHCs, requires some form of spatial coherence (***Zweig and Shera, 1995***; ***Talmadge et al., 1998***; ***Shera and Guinan, 2007***). Since incoherent noise sources do not satisfy these conditions, the criticism raised by ***van der Heijden and Vavakou (2021)***—namely, that that the responses of active models inevitably succumb to the catastrophic “cascaded” amplification of distributed, incoherent noise sources—appears puzzling.

5 In their recent critique of the cochlear amplifier, ***van der Heijden and Vavakou (2021)*** claim that their own theoretical speculations differ from most models in that they focus on the abrupt transition between long- and short-wave regions. In particular, the authors interpret the abrupt transition as the locus where cochlear waves convert from propagating without substantial loss to a mode where energy dissipates via mechanical vibrations within the sensory epithelium. Their criticisms, however, mischaracterize existing theories and appear at odd with the modeling literature (see e.g., ***de Boer, 1981***, ***1983***; ***Shera et al., 2005***; ***Zweig, 2015***; ***Altoè and Shera, 2020a***). Indeed, we have recently demonstrated that the long-to-short wave transition region found in active models is abrupt enough that one can obtain good approximate solutions to the model equations using a “shotgun wedding” that matches long- and short-wave solutions at a seam (***Altoè and Shera, 2020b***).

6 Simple, hand-waving arguments might reassure the skeptical reader that near-CF BM displacement can reasonably be expected to decrease with increasing frequency. The acoustic power carried by the traveling wave is proportional to the fluid velocity (necessary equal to BM velocity near the BM), and the power delivered from the middle ear to the cochlea is largely independent of frequency (***Lynch et al., 1982***; ***Shera and Zweig, 1991***). Therefore, an approximately constant acoustic power entering the cochlea can be expected to produce approximately constant BM velocity at the peak of the traveling wave, even for a passive cochlea where there is no OHC amplification (***Zweig et al., 1976***). Constant velocity, however, implies that the corresponding displacement decreases as 1∕*f*, consistent with the data in Fig. 2. Thus, the observed scaling with frequency is neither a dubious assumption of byzantine models or an inexplicable peculiarity of the murine cochlea.

7 Cascading *N* low-pass filters and mixing their responses (i.e., computing a weighted sum) is equivalent to producing a filter with *N* − 1 zeros and *N* poles, where the pole locations are determined by the poles of the individual filters and the zeros depend on the mixing coefficients. Mixing the result with the input signal increases the number of zeros to *N*. When feedback loops are included, the system of cascaded low-pass filters becomes a general purpose, zero-pole model of order *N*. Such models are commonly used in system identification, signal processing, and physical modeling applications (e.g., ***Choi, 2012***).

## Notes

### Competing Interest Statement

The authors have declared no competing interest.

## References

Allen JB. Nonlinear cochlear signal processing. In: Physiology of the Ear, Second Edition Singular Thompson; 2001.p. 393–442.

Altoè A, Shera CA. The cochlear ear horn: Geometric origin of tonotopic variations in auditory signal processing. Sci Rep. 2020; 10:20528.

Altoè A, Charaziak KK, Shera CA. Dynamics of cochlear nonlinearity: Automatic gain control or instantaneous damping? J Acoust Soc Am. 2017; 142:3510–3519.

Altoè A, Pulkki V, Verhulst S. The effects of the activation of the inner-hair-cell basolateral K+ channels on auditory nerve responses. Hear Res. 2018; 364:68–80.

Altoè A, Shera CA. Nonlinear cochlear mechanics without direct vibration-amplification feedback. Phys Rev Res. 2020; 2:013218.

Ashmore JF. A fast motile response in guinea-pig outer hair cells: The cellular basis of the cochlear amplifier. J Physiol. 1987; 388:323–347.

Ashmore JF. Cochlear outer hair cell motility. Physiol Rev. 2008; 88:173–210.

Beurg M, Evans MG, Hackney CM, Fettiplace R. A large-conductance calcium-selective mechanotransducer channel in mammalian cochlear hair cells. J Neurosci. 2006; 26:10992–11000.

Brownell WE, Bader CR, Bertrand D, De Ribaupierre Y. Evoked mechanical responses of isolated cochlear outer hair cells. Science. 1985; 227:194–196.

Caprara GA, Mecca AA, Wang Y, Ricci AJ, Peng AW. Hair bundle stimulation mode modifies manifestations of mechanotransduction adaptation. J Neurosci. 2019; 39:9098–9106.

Choi B. ARMA model identification. Springer Science & Business Media; 2012.

Cody AR, Russell IJ. The responses of hair cells in the basal turn of the guinea-pig cochlea to tones. J Physiol. 1987; 383:551–569.

Dallos P, Evans BN. High-frequency motility of outer hair cells and the cochlear amplifier. Science. 1995; 267:2006–2009.

Dallos P, Wu X, Cheatham MA, Gao J, Zheng J, Anderson CT, Jia S, Wang X, Cheng WH, Sengupta S, et al. Prestin-based outer hair cell motility is necessary for mammalian cochlear amplification. Neuron. 2008; 58:333–339.

de Boer E. Short waves in three-dimensional cochlea models: Solution for a ‘block’ model. Hear Res. 1981; 4:53–77.

de Boer E. No sharpening? A challenge for cochlear mechanics. J Acoust Soc Am. 1983; 73:567–573.

Dewey JB, Altoè A, Shera CA, Applegate BE, Oghalai JS. Cochlear outer-hair-cell electromotility enhances organ-of-Corti motion on a cycle-by-cycle basis at high frequencies. Proc Natl Acad Sci USA. 2021; 118:e2025206118.

Dewey JB, Applegate BE, Oghalai JS. Amplification and suppression of traveling waves along the mouse organ of Corti: Evidence for spatial variation in the longitudinal coupling of outer hair cell-generated forces. J Neurosci. 2019; 39:1805–1816.

Dong W, Olson ES. Detection of cochlear amplification and its activation. Biophys J. 2013; 105:1067–1078.

Frank G, Hemmert W, Gummer AW. Limiting dynamics of high-frequency electromechanical transduction of outer hair cells. Proc Natl Acad Sci USA. 1999; 96:4420–4425.

Guinan JJ. How are inner hair cells stimulated? Evidence for multiple mechanical drives. Hear Res. 2012; 292:35–50.

He DZ, Jia S, Dallos P. Mechanoelectrical transduction of adult outer hair cells studied in a gerbil hemicochlea. Nature. 2004; 429:766–770.

He W, Ren T. The origin of mechanical harmonic distortion within the organ of Corti in living gerbil cochleae. Commun Biol. 2021; 4:1–11.

van der Heijden M, Vavakou A. Rectifying and sluggish: outer hair cells as regulators rather than amplifiers. Hear Res. 2021; p. 108367.

Housley GD, Ashmore JF. Ionic currents of outer hair cells isolated from the guinea-pig cochlea. J Physiol. 1992; 448:73–98.

Iwasa KH. Negative membrane capacitance of outer hair cells: Electromechanical coupling near resonance. Sci Rep. 2017; 7:12118.

Johnson SL, Beurg M, Marcotti W, Fettiplace R. Prestin-driven cochlear amplification is not limited by the outer hair cell membrane time constant. Neuron. 2011; 70:1143–1154.

Kemp DT. Stimulated acoustic emissions from within the human auditory system. J Acoust Soc Am. 1978; 64:1386–1391.

Koch C. Biophysics of computation: information processing in single neurons. Oxford university press; 2004.

Kolston PJ. The importance of phase data and model dimensionality to cochlear mechanics. Hear Res. 2000; 145:25–36.

Kolston PJ, de Boer E, Viergever MA, Smoorenburg GF. What type of force does the cochlear amplifier produce? J Acoust Soc Am. 1990; 88:1794–1801.

Kros C, Crawford A. Potassium currents in inner hair cells isolated from the guinea-pig cochlea. The Journal of Physiology. 1990; 421:263–291.

Lee HY, Raphael PD, Xia A, Kim J, Grillet N, Applegate BE, Bowden AKE, Oghalai JS. Two-dimensional cochlear micromechanics measured in vivo demonstrate radial tuning within the mouse organ of Corti. J Neurosci. 2016; 36:8160–8173.

Liberman MC, Gao J, He DZZ, Wu X, Jia S, Zuo J. Prestin is required for electromotility of the outer hair cell and for the cochlear amplifier. Nature. 2002; 419:300–304.

Lighthill J. Energy flow in the cochlea. J Fluid Mech. 1981; 106:149–213.

Liu YW, Neely ST. Outer hair cell electromechanical properties in a nonlinear piezoelectric model. J Acoust Soc Am. 2009; 126:751–761.

Lu TK, Zhak S, Dallos P, Sarpeshkar R. Fast cochlear amplification with slow outer hair cells. Hear Res. 2006; 214:45–67.

Lynch TJ, Nedzelnitsky V, Peake WT. Input impedance of the cochlea in cat. J Acoust Soc Am. 1982; 72:108–130.

Mountain DC, Hubbard AE. A piezoelectric model of outer hair cell function. J Acoust Soc Am. 1994; 95:350–354.

Nam H, Guinan JJ. Low-frequency bias tone suppression of auditory-nerve responses to low-level clicks and tones. Hear Res. 2016; 341:66–78.

Nobili R, Mammano F. Biophysics of the cochlea II: Stationary nonlinear phenomenology. J Acoust Soc Am. 1996; 99:2244–2255.

Nowotny M, Gummer AW. Nanomechanics of the subtectorial space caused by electromechanics of cochlear outer hair cells. Proc Natl Acad Sci USA. 2006; 103:2120–2125.

Ospeck M, Iwasa KH. How close should the outer hair cell RC roll-off frequency be to the characteristic frequency? Biophys J. 2012; 102:1767–1774.

Perez-Flores MC, Lee JH, Park S, Zhang XD, Sihn CR, Ledford HA, Wang W, Kim HJ, Timofeyev V, Yarov-Yarovoy V, et al. Cooperativity of Kv7. 4 channels confers ultrafast electromechanical sensitivity and emergent properties in cochlear outer hair cells. Sci Adv. 2020; 6:eaba1104.

Peterson AJ, Heil P. Phase locking of auditory-nerve fibers: The role of lowpass filtering by hair cells. J Neurosci. 2020; 40:4700–4714.

Peterson LC, Bogert BP. A dynamical theory of the cochlea. J Acoust Soc Am. 1950; 22:369–381.

Rabbitt RD. The cochlear outer hair cell speed paradox. Proc Natl Acad Sci USA. 2020; 117:21880–21888.

Ren T, He W, Barr-Gillespie PG. Reverse transduction measured in the living cochlea by low-coherence hetero-dyne interferometry. Nat Comm. 2016; 7.

Ren T, He W, Kemp DT. Reticular lamina and basilar membrane vibrations in living mouse cochleae. Proc Natl Acad Sci USA. 2016; 113:9910–9915.

Russell IJ, Cody AR, Richardson GP. The responses of inner and outer hair cells in the basal turn of the guinea-pig cochlea and in the mouse cochlea grown in vitro. Hear Res. 1986; 22:199–216.

Santos-Sacchi J. On the frequency limit and phase of outer hair cell motility: Effects of the membrane filter. J Neurosci. 1992; 12:1906–1916.

Santos-Sacchi J, Dilger JP. Whole cell currents and mechanical responses of isolated outer hair cells. Hear Res. 1988; 35:143–150.

Santos-Sacchi J, Tan W. The frequency response of outer hair cell voltage-dependent motility is limited by kinetics of prestin. J Neurosci. 2018; 38:5495–5506.

Sasmal A, Grosh K. Unified cochlear model for low-and high-frequency mammalian hearing. Proc Natl Acad Sci USA. 2019; 116:13983–13988.

Sasmal A, Grosh K. The competition between the noise and shear motion sensitivity of cochlear inner hair cell stereocilia. Biophys J. 2018; 114:474–483.

Scherer MP, Gummer AW. Impedance analysis of the organ of Corti with magnetically actuated probes. Biophys J. 2004; 87:1378–1391.

Shera CA, Guinan JJ. Cochlear traveling-wave amplification, suppression, and beamforming probed using non-invasive calibration of intracochlear distortion sources. J Acoust Soc Am. 2007; 121:1003–1016.

Shera CA. Intensity-invariance of fine time structure in basilar-membrane click responses: Implications for cochlear mechanics. J Acoust Soc Am. 2001; 110:332–348.

Shera CA. Laser amplification with a twist: Traveling-wave propagation and gain functions from throughout the cochlea. J Acoust Soc Am. 2007; 122:2738–2758.

Shera CA, Tubis A, Talmadge CL. Coherent reflection in a two-dimensional cochlea: Short-wave versus long-wave scattering in the generation of reflection-source otoacoustic emissions. J Acoust Soc Am. 2005; 118:287–313.

Shera CA, Zweig G. A symmetry suppresses the cochlear catastrophe. J Acoust Soc Am. 1991; 89:1276–1289.

Shera CA, Zweig G. An empirical bound on the compressibility of the cochlea. J Acoust Soc Am. 1992; 92:1382–1388.

Soons JA, Ricci AJ, Steele CR, Puria S. Cytoarchitecture of the mouse organ of Corti from base to apex, determined using in situ two-photon imaging. J Assoc Res Otolaryngol. 2015; 16:47–66.

Taberner AM, Liberman MC. Response properties of single auditory nerve fibers in the mouse. J Neurophysiol. 2005; 93:557–569.

Talmadge CL, Tubis A, Long GR, Piskorski P. Modeling otoacoustic emission and hearing threshold fine structures. J Acoust Soc Am. 1998; 104:1517–1543.

Vavakou A, Cooper NP, van der Heijden M. The frequency limit of outer hair cell motility measured in vivo. Elife. 2019; 8:e47667.

Zheng J, Shen W, He DZ, Long KB, Madison LD, Dallos P. Prestin is the motor protein of cochlear outer hair cells. Nature. 2000; 405:149–155.

Zweig G. Finding the impedance of the organ of Corti. J Acoust Soc Am. 1991; 89:1229–1254.

Zweig G. Linear cochlear mechanics. J Acoust Soc Am. 2015; 138:1102–1121.

Zweig G. Nonlinear cochlear mechanics. J Acoust Soc Am. 2016; 139:2561–2578.

Zweig G, Lipes R, Pierce JR. The cochlear compromise. J Acoust Soc Am. 1976; 59:975–982.

Zweig G, Shera CA. The origin of periodicity in the spectrum of evoked otoacoustic emissions. J Acoust Soc Am. 1995; 98:2018–2047.

